# Recurrent neural circuits overcome partial inactivation by compensation and re-learning

**DOI:** 10.1101/2021.11.12.468273

**Authors:** Colin Bredenberg, Cristina Savin, Roozbeh Kiani

## Abstract

Technical advances in artificial manipulation of neural activity have precipitated a surge in studying the causal contribution of brain circuits to cognition and behavior. However, complexities of neural circuits challenge interpretation of experimental results, necessitating theoretical frameworks for systematic explorations. Here, we take a step in this direction, using, as a testbed, recurrent neural networks trained to perform a perceptual decision. We show that understanding the computations implemented by network dynamics enables predicting the magnitude of perturbation effects based on changes in the network’s phase plane. Inactivation effects are weaker for distributed network architectures, are more easily discovered with non-discrete behavioral readouts (e.g., reaction times), and vary considerably across multiple tasks implemented by the same circuit. Finally, networks that can “learn” during inactivation recover function quickly, often much faster than the original training time. Our framework explains past empirical observations by clarifying how complex circuits compensate and adapt to perturbations.

## 2 Introduction

Artificial manipulations of neural circuits are vital tools in modern neuroscience for investigating the neural computations that underlie behavior. These manipulation techniques include lesions (Vaidya et al. 2019, Newsome & Pare 1988), pharmacological inactivation (Katz et al. 2016, Hanks et al. 2015), microstimulation (Salzman et al. 1990), optogenetic (Fetsch et al. 2018, Brown et al. 2018, Tremblay et al. 2020, Rajasethupathy et al. 2016), and chemogenetic (Tervo et al. 2014, Eldridge et al. 2016) inactivation or excitation. It is commonly assumed that the results of these experiments are easily interpretable, i.e., that changes of behavior following artificial inactivation or excitation of a circuit demonstrate the importance of that circuit in producing that behavior. However, the converse of this statement—that the absence of changes in behavior following circuit manipulation indicate that this circuit does not play a role in producing the behavior—is much more difficult to assert.

Beyond sensory bottlenecks, the distributed nature of computations for higher brain functions makes attribution of a single function to a single circuit much more challenging, because multiple areas may jointly contribute to a function, and may be able to mutually compensate for inactivity in other re-gions (Wolff & OÖlveczky 2018). Furthermore, the capacity of neural circuits to compensate for inactivation is well documented (Vaidya et al. 2019). This implies that other neurons in the circuit or areas of the brain which are not normally causal in producing a particular behavior can adapt to play an important role. Transient manipulation may produce effects that are more difficult to adapt to, but there is evidence for compensation for even transient optogenetic inactivation (Fetsch et al. 2018).

Since the effect of perturbations may not always follow simple intuition, modelling can provide a useful way to reason about possible experimental outcomes and their interpretation. For this we turn to artificial recurrent neural networks (RNNs), which have been successfully used as a bridge between neural activity and behavior in several tasks (Rigotti et al. 2010, Mante et al. 2013, Yang et al. 2019). RNNs are powerful model systems that share many complexities of brain circuits, while permitting direct access to the inner-workings of the system. Critically, the choice of architecture and training objectives allows the exploration of different circuit scenarios, where the contribution of network elements to the output is known. Moreover, simulations provide the ability of perturbing activity with any spatial and temporal resolution. The combination of these features results in immense control and knowledge about complex networks, at a level unattainable in real brain circuits. We build on this foundation to investigate the ability of causal interventions to reveal the role neurons and populations play in the distributed computations performed in a complex network.

As a specific example of the complexities involved in causal manipulation of neural circuits, we focus on the integration of sensory evidence for the random dots motion (RDM) task, a decision making paradigm that engages multiple frontoparietal cortices (Kiani, Cueva, Reppas & Newsome 2014, Roitman & Shadlen 2002, Kim & Shadlen 1999, Mochol et al. 2021) and subcortical areas (Ding & Gold 2013, Horwitz & Newsome 1999, Ratcliff et al. 2011), and which has been a target for many causal studies, with contradictory outcomes (Katz et al. 2016, Zhou & Freedman 2019*a*, Hanks et al. 2006, Licata et al. 2017, Erlich et al. 2015). These studies suggest a distributed process for neural implementation of decision-making, whose complexity challenges standard experimental techniques for identifying causal relationships between individual brain regions and behavior. On the modeling side, several successful RNN-based models have been used to replicate neural response dynamics and behavior in perceptual decision making (Mante et al. 2013, Wong & Wang 2006, Rigotti et al. 2010) and their dynamical systems properties are well understood. In particular, recurrent networks can perform near-optimal evidence integration by constructing a low-dimensional attractor (Goldman et al. 2003, Wong & Wang 2006, Cain et al. 2013), whose structure provides a direct route to investigating the computational integrity of the circuit.

Here, we use similar trained RNN circuits to systematically study causal manipulations of the RDM task. We show that inactivation of subsets of neurons in the network affects behavior by damaging the low-dimensional attractor, with larger activity perturbations having a greater impact on both accuracy and reaction time, more so for the latter. In a more complex network, where integration is done collaboratively by multiple circuits, inactivation may or may not affect the computational structure of the solution and the behavior. In particular, in networks with parallel, redundant computation, inactivation of a subset of the circuit can have little to no effect on the network output. We further show that in networks trained to perform multiple tasks, inactivation can have surprisingly inconsistent effects across tasks. It may affect one task and not the other, even though the circuit is designed to be necessary for both. Lastly, we demonstrate that recurrent neural networks that retain plasticity — and continue learning — after the inactivation reconfigure themselves to regain the accuracy they had prior to inactivation. The speed of recovery depends on the extent and temporal profile of the inactivation, and network performance is closely related to the integrity of the network’s attractors. These observations caution against simplistic interpretations of causal experiments and suggest concrete ways to avoid interpretational pitfalls and improve experimental design.

## 3 Results

### 3.1 Hierarchical recurrent networks approximate linear integration for simple sensory decisions

We begin with the simplest hierarchical recurrent network architecture (Fig. 1a) consisting of a sensory-like population (P1) and an integrating population (P2). Neurons in each population have dense recurrent connections between them, while the sensory population projects sparsely to the integrating population. The P2 population roughly corresponds to the collection of the recurrently connected cortical and sub-cortical neurons involved in the decision-making process; however, it does not reflect the precise anatomy of brain networks. The stimulus in each trial randomly fluctuates around a mean stimulus strength level, akin to the dynamic random dots stimuli in direction discrimination tasks (Newsome et al. 1989, Roitman & Shadlen 2002) where motion energy fluctuates around a mean dictated by the coherence of random dots. This fluctuating stimulus input is received by the sensory population P1, and relayed to the integrating population P2. The network is trained such that a linear read-out of the activity of population P2 at each moment matches the integral of the stimulus input up to that time. All connections in the network are plastic during training and modified by backpropagation through time (BPTT) (see Methods). After learning, the sensory population shows coherence tuning (see example response profiles in Fig. S1a-c), while the integration population develops response profiles — ramping activity — similar to those reported in posterior parietal and prefrontal cortex (Fig. S1d-f). The connections are fixed after the initial training, reflecting the common assumption that synapses and network dynamics do not significantly change after inactivation. We separately also explore the effects of continuous plasticity on causal experiments (section 3.6).

**Figure 1:**
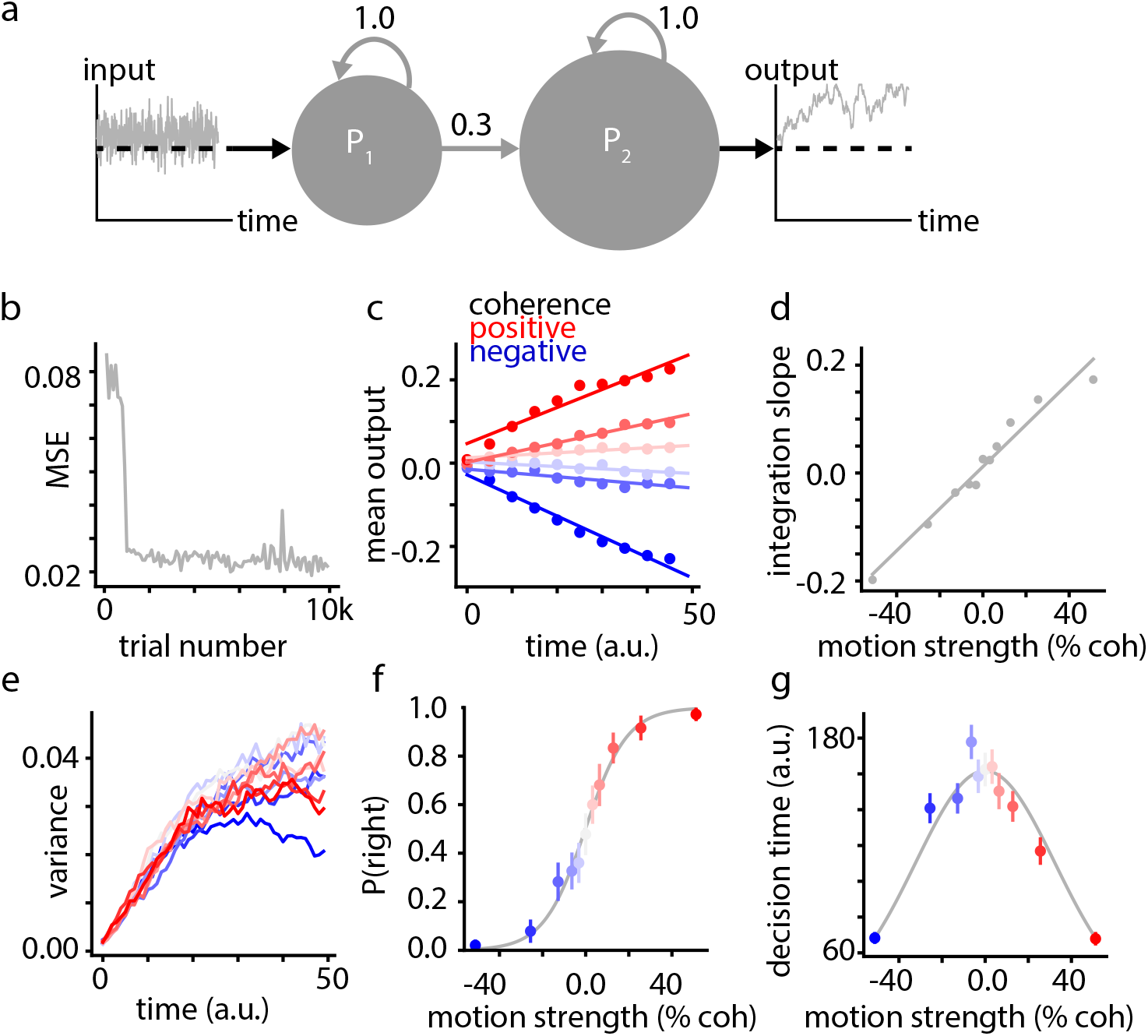
A two-stage hierarchical RNN performing linear integration of noisy inputs for a sensory decision-making task. **a)** Network schematic. **b)** Network learning throughout time in units of mean-squared error. **c)** Mean activity of the output unit after training for different stimulus strengths (motion coherence). Model outputs (solid points) increase linearly over time up to a saturating level implemented in the training procedure (see Methods). Lines are fits to the data points over the time range [0 50], measured in arbitrary network time units. **d)** The slope of changes in model output as a function of stimulus strength. **e)** Variance of model output as a function of time for different stimulus strengths. The linear increase is expected for integration of noisy inputs over time. The late saturation and decline of the variance, especially for stronger stimuli, is caused by the bound on the integration process. **f)** Psychometric function of a trained model. Data points show the probability of choosing right for different stimulus strengths ranging from strong leftward motion (negative coh) to strong rightward (positive coh). The gray curve is a logistic fit. Error bars show ± 1 s.e.m. **g)** Chronometric function of a trained model. Data points show the mean time the model output takes to reach the decision bounds (decision time). The gray curve is a Gaussian function fit to the data points.

The network is trained within a couple of thousands of trials (Fig. 1b), similar to training schedules for nonhuman primates (Gold et al. 2010). After training, the P2 output closely approximates the integral of the stimulus input over time. Two hallmarks of temporal integration are linear scaling with time (Fig. 1c) and stimulus strength (Fig. 1c-d) (Gold & Shadlen 2007): if a network is receiving a constant-mean stimulus input, the integrated output will be a linear function over time, with slope equal to the mean of the stimulus. The model output represents linear integration for a wide range of inputs and times but saturates for very large values of the integral. Because of the limited dynamic range of the neural responses, the curtailed range of the integral improves the precision of the representation of the integrated evidence for weaker stimuli, where the network precision matters the most for the accuracy of choices. Another hallmark of temporal integration of noisy inputs is linear growth of the variance of the integral over time. The motion energy of the random dots stimulus at each time is an independent sample from a normal distribution, so their sum over time–integration–should have a variance that scales linearly with time (Roitman & Shadlen 2002, Churchland et al. 2011). Our network output captures this hallmark of optimal integration (Fig. 1e).

Since the network’s integration process stops when the network output reaches a fixed decision bound, the model provides trial-by-trial estimates of the network’s decision time and choice. The time to bound is the decision time, and the sign of the network output at the time of bound crossing determines the choice (right or leftward motion). Our model decision times are in units of the network time steps. The resulting psychometric and chronometric functions of the model show profiles qualitatively similar to experimental results, in particular, faster, more accurate responses for stronger motion stimuli (Fig. 1f-g) (Roitman & Shadlen 2002, Kiani, Corthell & Shadlen 2014, Palmer et al. 2005).

### 3.2 Behavioral effects of inactivation grow with the size of the inactivated population

We explored the effects of inactivation on this circuit by selectively silencing a proportion of neurons in the integrating population (Fig. 2a), and analyzing the inactivation effects on the output of the model. For a particular trained network, we measured the change in the psychometric and chronometric functions after perturbation as a means to characterize the effects of inactivation. We found that decision times are strongly sensitive to inactivation. Weak inactivations (5-10% of the population) moderately increase the decision time of the network, and medium and strong perturbations (20% and 30% of the population, respectively) cause a much larger increase (Fig. 2d,g).

**Figure 2:**
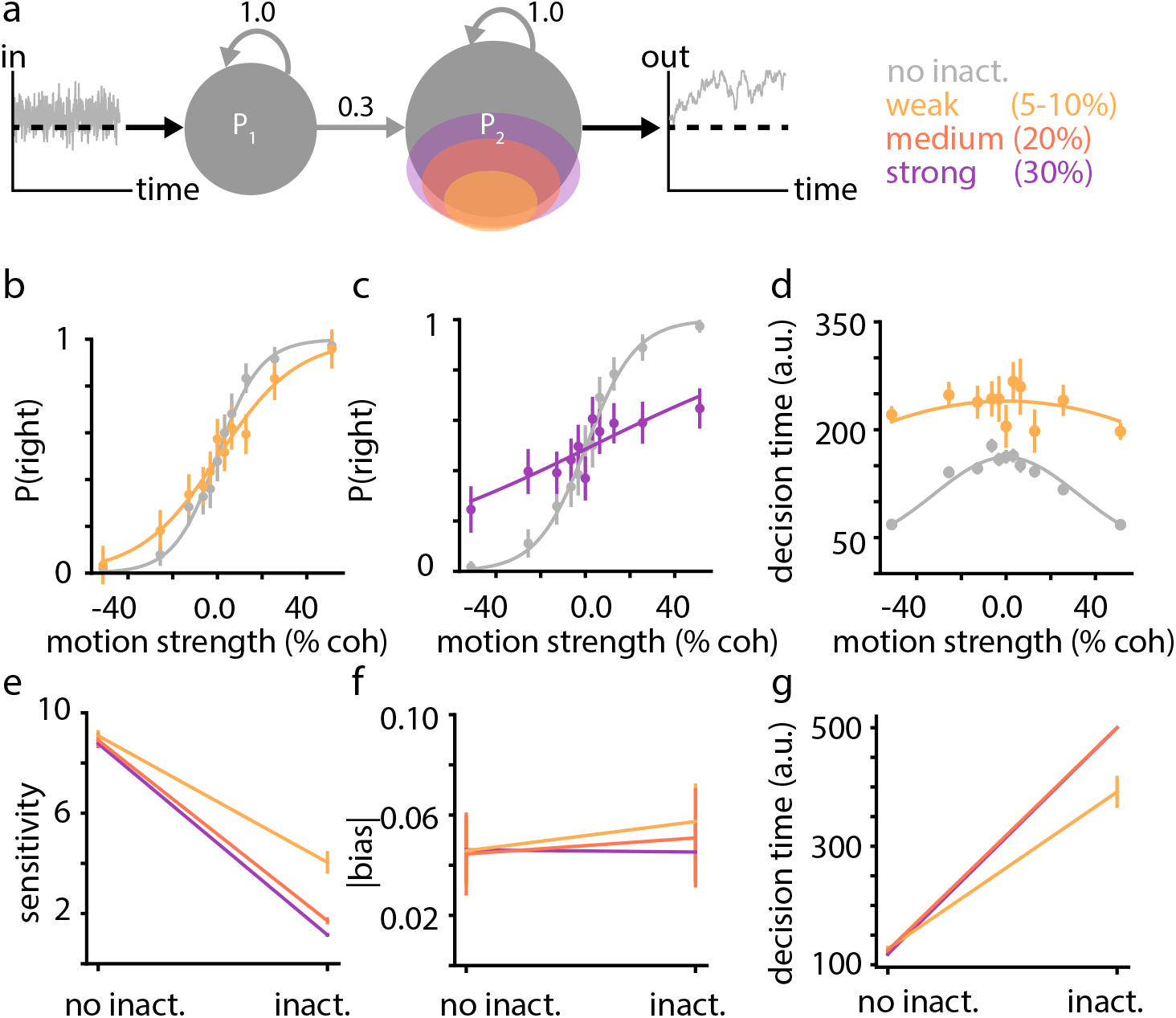
Inactivation of the integrating circuit reduces sensitivity and increases decision times, with larger effects when a larger portion of neurons are silenced. **a)** Inactivation schematic. Colored ovals indicate the affected proportion of the P2 network. **b-c)** Psychometric functions of example networks after training (grey) and after a weak (5-10%, b) or medium-size (20%, c) inactivation. **d** Chronometric function of the same network in panel b following weak inactivation. **e)** Changes of sensitivity (slope of psychometric function) across 10 trained networks for various inactivation sizes. **f)** Effects of inactivation on bias (shift of the mid-point of the psychometric function from 0% coherence) across trained networks. **g)** Changes of mean decision times across trained networks. Error bars show s.e.m. Maximal trial duration set to 500 steps.

The effect of perturbation on choice was more variable and complex. We quantified these effects by extracting measures of the sensitivity and bias for the psychometric functions, and calculating the change in these measures with weak, medium, and strong inactivation. The sensitivity of the psychometric function decreased as more of the neurons were affected, with a corresponding decrease in the average sensitivity with inactivation size (Fig. 2b-c,e). The magnitude of the bias, however, minimally changed across inactivation levels (Fig. 2f), suggesting that the primary loss of function caused by increasing the perturbation magnitude is a loss of sensitivity. Overall, in our basic network architecture, even weak perturbations decreased sensitivity and substantially increased reaction time, with the magnitude of these effects increasing with the magnitude of inactivation.

### 3.3 Inactivation effects arise from perturbing the structure of the underlying population dynamics

The optimal solution for random dots motion discrimination involves integration along a 1-dimensional axis (Wald & Wolfowitz 1950, Shadlen et al. 2006, Drugowitsch et al. 2012, Khalvati et al. 2021), so the dynamics of the trained network are likely to lie on a low dimensional manifold (Ganguli & Sompolinsky 2012). Indeed, simple dimensionality reduction using PCA shows that the circuit dynamics are approximately one dimensional, with the first principal axis explaining about 70% of the neural response variance (Fig. 3a). The low dimensional structure of the neural activity allows us to project the full network dynamics on the first principal component axis. In this space we can mathematically analyze the dynamical features of the trained network that enable it to perform evidence integration.

**Figure 3:**
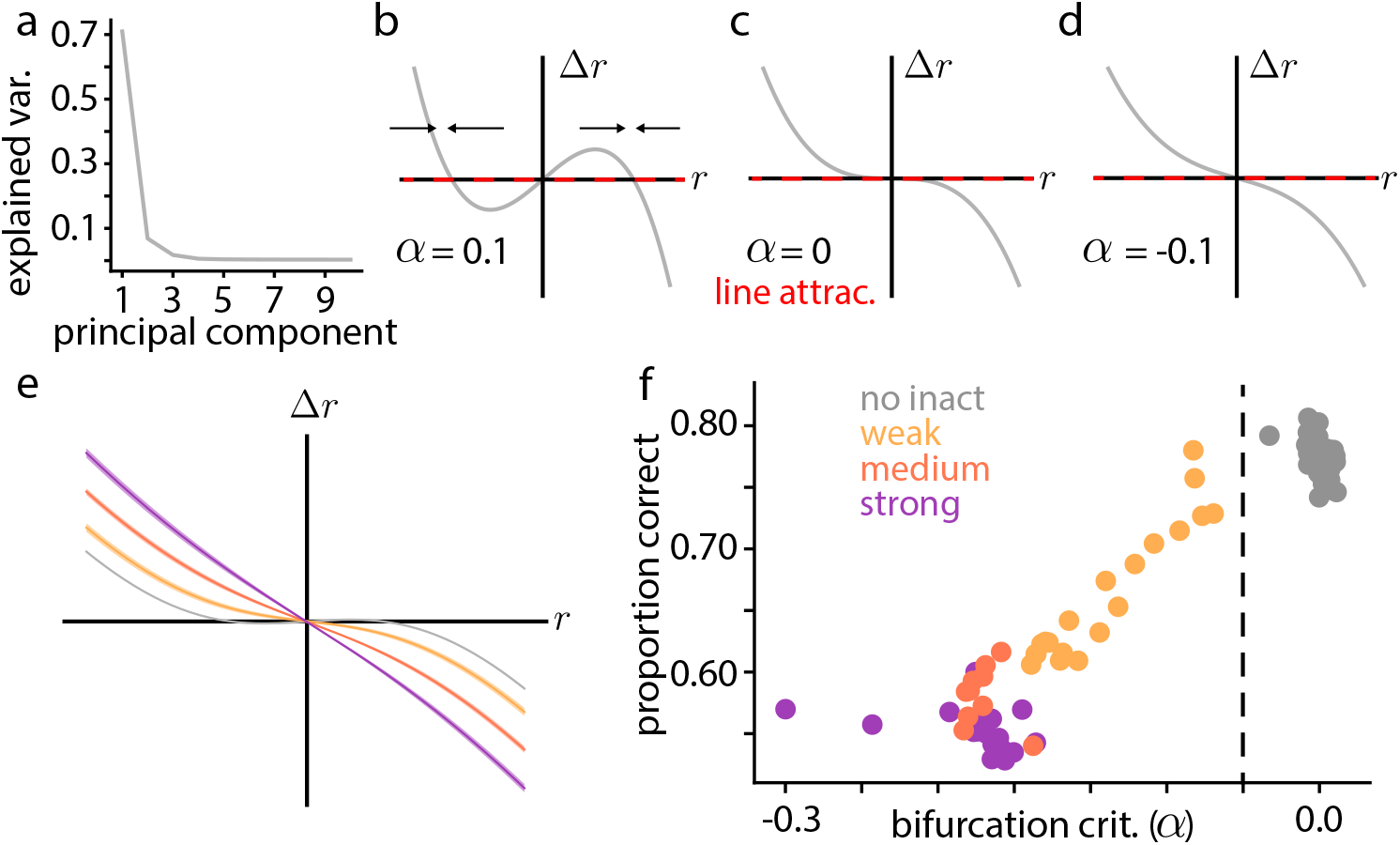
Integrating network implements a shallow bistable attractor, whose disruption determines the magnitude of the behavioral effects of inactivation. **a)** Fraction of explained variance as a function of the number of latent dimensions of the network responses to test stimuli. **b-d)** Schematic of a pitchfork bifurcation. The red dashed line shows the phase portrait for a line attractor implementing optimal evidence integration. For *α* > 0, the network has two stable attractors (b, arrows indicate sign of Δ*r*), for *α* = 0, a saddle point (c), and for *α* < 0, a single stable attractor (d). **e)** Phase plot for the reduced network before (gray) and after (colors) perturbation. Shaded regions indicate s.e.m. across network realizations. **f)** Fraction of correct responses as a function of the bifurcation criterion estimated for each network.

In the absence of a stimulus, the network activity can be approximated as one-dimensional population dynamics of the general functional form:

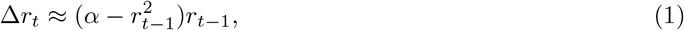

where Δ*r*_*t*_ is the change in population activity across time and the value of parameter *α* and the constant of proportionality depend on the trained network weights (See Methods). Different settings of *α* change the dynamical properties of the system and its ability to solve the evidence integration task. This property is illustrated in the phase plane in Figure 3b, which describes the relationship between Δ*r*_*t*_ and *r*_*t−*1_. When *α* is positive, the dynamics exhibit three fixed points (corresponding to values of *r*_*t*_ for which Δ*r*_*t*_ is zero; Fig. 3b). Two of these are attracting, separated by an unstable fixed point at *r*_*t*_ = 0: when starting from a positive value of activity *r*, the network will eventually converge to the positive fixed point, and similarly a negative starting condition will converge to the negative fixed point. Sensory drive to the network will push the dynamics towards one or the other, eventually converging to a final binary decision. This is similar to the phase plane of previous circuit models of evidence integration based on bistable attractor dynamics (Wong & Wang 2006).

Ideal evidence integration sums all incoming inputs, 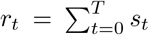,, and follows slightly different dynamics. In the phase plane, ideal integration means that *r*_*t*_ changes at every time step as

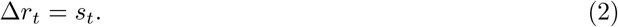

Hence, the ideal solution for evidence integration involves a line attractor, where no change in activity occurs in the absence of input (Δ*r*_*t*_ = 0 whenever *s*_*t*_ = 0; red line in Fig. 3b-d). The trained RNN approximates this solution when *α* is close to zero (Fig. 3c). Once *α* becomes negative, the network will start to behave qualitatively differently (formally, this corresponds to a pitchfork bifurcation (Strogatz 2018), which is why we refer to *α* as the bifurcation criterion). In this regime, the network has only one stable fixed point at the origin (Fig. 3d). Changes in network activity caused by new stimuli rapidly relax to this fixed point, causing the network to lose its ability to integrate.

The value of the bifurcation criterion *α*, which we derive directly from the RNN activity, captures the key dynamic properties of the model networks and predicts a given network’s ability to perform evidence integration. Indeed, trained networks generally have a small positive *α*, corresponding to a shallow bi-stable attractor and close-to-ideal evidence integration (Fig. 3e).

Different forms of causal interventions, such as inactivation, will result in altered population dynamics, with a corresponding change in the bifurcation criterion *α*. In particular, our inactivation experiments push the network past the bifurcation point: as the magnitude of the inactivation increases, *α* values become increasingly negative (Fig. 3e,f). The remaining fixed point leads to forgetting past inputs and correspondingly poor performance (Fig. 3f).

Overall, these results establish that our network approximates integration within a bounded region of state space via a shallow bi-stable attractor, and that the loss of function caused by perturbations is due to the loss of this attractor structure. This dynamical systems analysis paints a more refined picture of causal interventions in the random dots motion discrimination task: inactivations that disrupt the computational structure embedded in the network (i.e., the bi-stable attractor) will produce behavioral impairments, while those that leave the attractor unaffected will not. While in our simple network architecture (Fig. 2a) all interventions disrupt the attractor structure, this may not necessarily be the case for more complex distributed networks, as we will see below.

### 3.4 In distributed architectures, inactivation effects can be variable

Exploring the effects of inactivation in a unitary circuit performing integration reveals a qualitatively similar picture across network and effect sizes. In a mammalian brain, however, sensory decisions are enabled by a distributed network consisting of multiple interacting circuits (Shadlen & Kiani 2013, Waskom et al. 2019). Past electrophysiological studies have found neurons that represent integration of sensory evidence in the parietal cortex (Shadlen & Newsome 2001, Churchland et al. 2008), lateral frontal cortex (Kim & Shadlen 1999, Mante et al. 2013, Mochol et al. 2021), motor and premotor cortices (Peixoto et al. 2021, Chandrasekaran et al. 2017, Hanks et al. 2015, Thura & Cisek 2014), the basal ganglia (Ding & Gold 2013, Yartsev et al. 2018), superior colliculus (Horwitz & Newsome 1999, Basso et al. 2021), and cerebellum (Deverett et al. 2018). The distributed nature of the computation, paired with potential circuit redundancies, make inactivation studies difficult to interpret. This is especially true when inactivation of a subcircuit in the network fails to produce measurable changes of behavior. Other nodes of the network could change their activity in responses to the inactivation, compensating for its effects (Li et al. 2016). Furthermore, there are a variety of more complex scenarios compatible with negative results (Yoshihara & Yoshihara 2018, Jonas & Kording 2017, Murray & Baxter 2006, Dunn 2003).

Although a detailed exploration of the distributed network that underlies decisions in the brain is beyond the scope of this paper, we take a first step in assessing the effects of architecture on inactivation experiments. In particular, we replace the unitary network structure analyzed above with a parallel architecture, where sensory inputs drive the responses of two non-interacting populations that collectively shape the network output (Fig. 4a). We train this parallel network to perform the same sensory integration task, followed by inactivating all of the neurons in one of the two parallel nodes and assessing the behavioral outcomes of the manipulation across a range of network instances. We find that even in this minimal version of a distributed computation the effects of inactivation can be quite variable in terms of performance.

**Figure 4:**
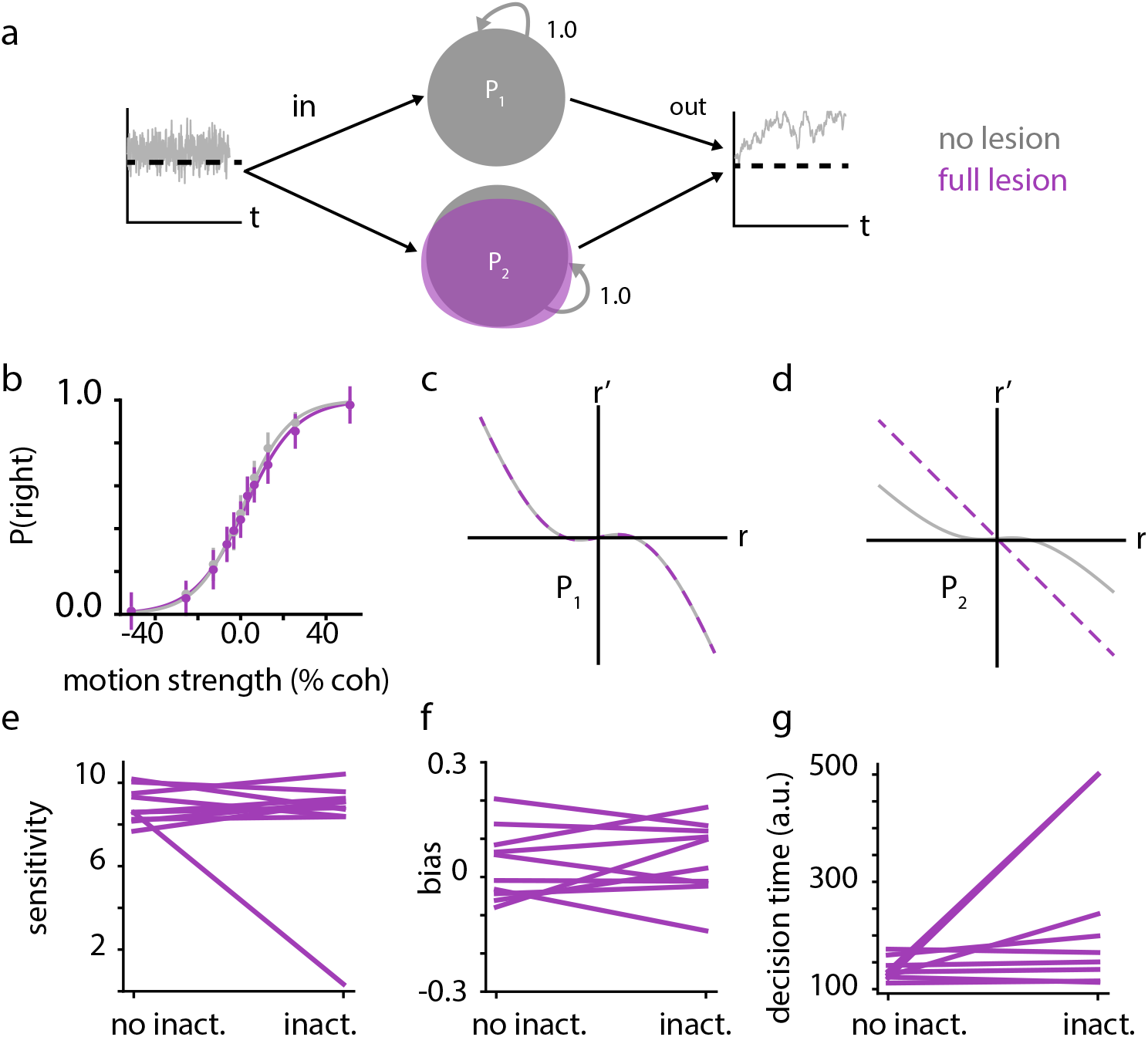
Distributing integration across multiple network nodes makes it resilient to disruptions in any single node. **a)** Schematic for a network with two parallel nodes (circuits) for integration computation. **b)** Psychometric curve for the parallel node network before and after a strong inactivation. The lines are logistic fits to the choices. Errors bars are s.e.m. **c)** Approximate phase portrait for the intact node, indicating a shallow bistable attractor. **d)** Phase portrait for the inactivated node. **e)** Sensitivity for the psychometric function before and after a strong inactivation. Each line shows an instance of the parallel node network with unique starting points and training history. Inactivation affected sensitivity in only one of the ten instances. **f)** Bias of the psychometric function before and after inactivation. **g)** Mean reaction time before and after inactivation.

Some networks exhibit minimal changes in the psychometric function due to inactivation (Fig. 4b, e), paired with a marked increase in reaction times (Fig. 4g). This phenomenology tracks back to the dynamical system properties of the underlying network. When examining the one-dimensional approximate phase portrait for each node in the network, we found that *both* exhibit the shallow bistable attractor dynamics indicative of approximately optimal sensory integration (Fig. 4c-d). The overall network output, which determines the final choice and decision time, is constructed by linearly combining the activities of both integrating populations. This architecture subsumes more specific architectures in which populations with distinct choice preference integrate evidence for their respective choices. The inactivation completely disrupts the attractor structure in the targeted sub-circuit P2 (Fig. 4d), but leaves the attractor in P1 intact (since they do not directly interact; Fig. 4c). Therefore, integration can still be performed using the intact sub-circuit. Nonetheless, the activity component from P2 is missing; as a result, the output could be weaker and it may take longer for the integrated signal to reach the same decision threshold, leading to slower responses. However, if the only measure of behavior is the choice, one may not notice any change of behavior, as evident in Fig 4.

A systematic investigation across networks with the same distributed architecture, but different trained connections, reveals that this inactivation-resistant solution is not universal: in some networks the sensitivity is largely unaffected, while others display a marked loss in sensitivity after inactivation (Fig. 4e). This variability traces back to the attractor structure of the individual solution found via learning. To solve the task, it is not strictly necessary to develop attractor structure in both circuits; a single attractor can do the job just as well, with the second circuit not truly involved in the computation. The effects of inactivation on the network that uses such an essentially unitary solution match natural intuitions: disrupting performance (sensitivity, bias) indicates that the sub-circuit is in fact involved in the network computation, while inactivating the sub-circuit that does not have attractor structure leaves the output essentially unaffected. However, once any real computational redundancy is in place, ‘negative results’ at the level of behavior need to be interpreted with caution. Absence of change in choice behavior following inactivation is insufficient to conclude that a certain network node lacks a functional role in the task.

Overall, this analysis reveals a more complicated picture of inactivation: disabling an individual node in a network will produce a loss of function only if no other node in the network is capable of compensating for its loss. Moreover, in dynamical systems such as the RNNs we study here, redundancy and compensation exist even in very simple networks, performing very simple tasks.

### 3.5 The effects of inactivation are task specific

Many perturbation experiments include inactivation of a circuit in more than one task, often to establish some form of dissociation in the involvement of the circuit across tasks. For example, inactivation of the lateral intraparietal cortex causes strong biases in a free choice task, where monkeys arbitrarily choose one of two targets that yield similar reward. But the same inactivation causes weaker or transient biases in perceptual tasks in which the choice is based on sensory stimuli (Katz et al. 2016, Zhou & Freedman 2019b, Jeurissen et al. 2021). Such dissociations are often interpreted as the circuit being more involved in the computations necessary for one task than the other. But this interpretation is not unique, and somewhat simplistic, as the operations of a complex, nonlinear dynamic system (e.g., neural circuits) could substantially differ in the presence and absence of a perturbation (see Discussion).

Consider a circuit designed to implement multiple computations, abstracted as different patterns of population activity in different tasks (Yang et al. 2019). Given an inactivation that affects a subset of neurons, are all tasks affected equally? Are some tasks more robust to inactivation than others? Even though in this thought experiment, the unperturbed circuit is known to underlie all the computations, its different activity patterns, and the corresponding dynamical features that implement the computations, could vary in their degree of sensitivity to different circuit perturbations.

To approach this challenge more directly, we trained a recurrent network to flexibly integrate one of two inputs, depending on a “context” cue (Fig. 5a). After learning, the network was able to integrate the cued input and ignore the other by embedding two distinct attractors in its dynamics, one for each context (Fig. 5b). As the network operates in context ‘1’, the psychometric and chronometric functions for the appropriate input qualitatively match those seen for simple integration, while the same functions calculated according to the off-context stimulus show sensitivity near 0. Similarly, decision times show strong dependence on the strength of the on-context input, and no dependence on the off-context input (Fig. 5c-d). This indicates that network responses are negligibly affected by the off-context input. This phenomenology holds robustly across different network realizations (Fig. 5e). The vast majority of the trained networks show sensitivity only to the on-context input, and a small minority of the trained networks show mixed sensitivity to both inputs — a suboptimal solution. Furthermore, this context-dependent integration occurs without any evident bias (Fig. 5f). In short, the trained networks behave as expected.

**Figure 5:**
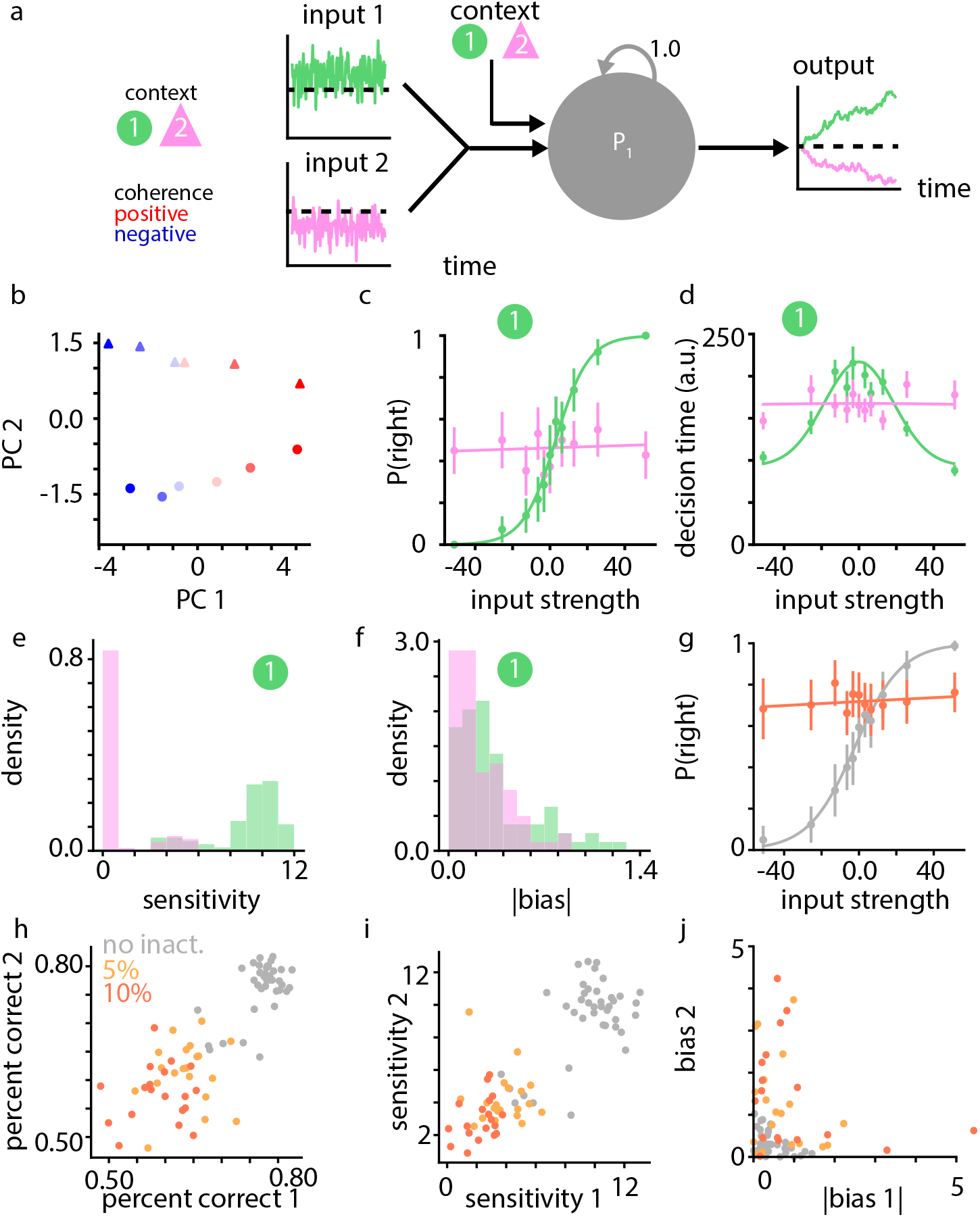
In a network trained to perform two similar computations (e.g., context-dependent integration of various inputs), effective inactivations influence performance in both tasks. **a)** Schematic of a network integrating either input 1 (green) or input 2 (pink) based on contextual inputs. **b**)Network steady-state activities projected onto the first two principal components. Color indicates the input mean (stimulus strength). Circles indicate context 1, and triangles indicate context 2. **c)** Psychometric function for responses in context 1 for input 1 (green) and input 2 (pink). The network successfully integrates the relevant input and ignores the irrelevant input based on context. **d)** Same as (c), but for the chronometric function. **e)** distribution of psychometric function sensitivities in context one for input 1 (green) and input 2 (pink). **f)** same as (e), but for the absolute value of the psychometric function bias. **g)** Psychometric function in context 1 pre-(grey) and post-inactivation (orange). **h)** Scatter plot of percent correct in context 1 vs. context 2. Grey corresponds to pre-inactivation, yellow to a 5% inactivation, and orange to a 10% inactivation. **i)** same as (h), but for psychometric function sensitivity. **j)** same as (h), but for psychometric function bias. Error bars indicate s.e.m.

When examining how inactivations affect performance on each task, an interesting picture emerges. Inactivating 10% of the neurons disrupted performance by decreasing the sensitivity of the network and by introducing a bias (Fig. 5g), in contrast to the previous simulations with a non-contextual integration task, where the primary loss-of-function was in the form of reduced sensitivity. Furthermore, different levels of inactivation resulted in correlated loss-of-function across contexts, in terms of both choice accuracy (Fig. 5h), and psychometric sensitivity (Fig. 5i). Biases increased for many inactivated networks (Fig. 5j), but appear to not increase jointly in both tasks. These results show that for this specific task (during context-specific integration), with inactivations of a random subset of neurons, loss-of-function appears to occur jointly across task contexts. This joint loss of function likely arises from the similarity of computations in the two tasks (integration) and the fact that we did not impose any training constraint to orthogonalize context-specific attractor dynamics. Depending on additional constraints, one could expect distinct results, possibly including independent changes of sensitivity in the two contexts.

A second example highlights distinct effects of inactivation across tasks. We trained our simple network architecture (Fig. 6a) to perform different computations in the two tasks, depending on context cues on each trial: integration of the stimulus, where the activity of the output neuron should match the time integral of the input, or a simple replication task, where the activity of the output neuron should match the network input (see Methods). We found that integration was more prone to be affected by inactivation. Across simulated networks with 5% and 10% inactivated neurons, all networks showed degraded sensitivity in the integration task, whereas the replication task showed a high degree of variability. Furthermore, when the replication task was affected by the inactivation, errors tended to be small overall compared to the input variance (normalized error before inactivation: 0.029 ± 0.002 std; after a 5% inactivation: 0.075 ± 0.053 std; and after a 10% inactivation: 0.093 ± 0.059 std). This result demonstrates that the presence of inactivation effect in one task and its absence in another task do not indicate that the circuit does not contribute to the computations in both tasks.

**Figure 6:**
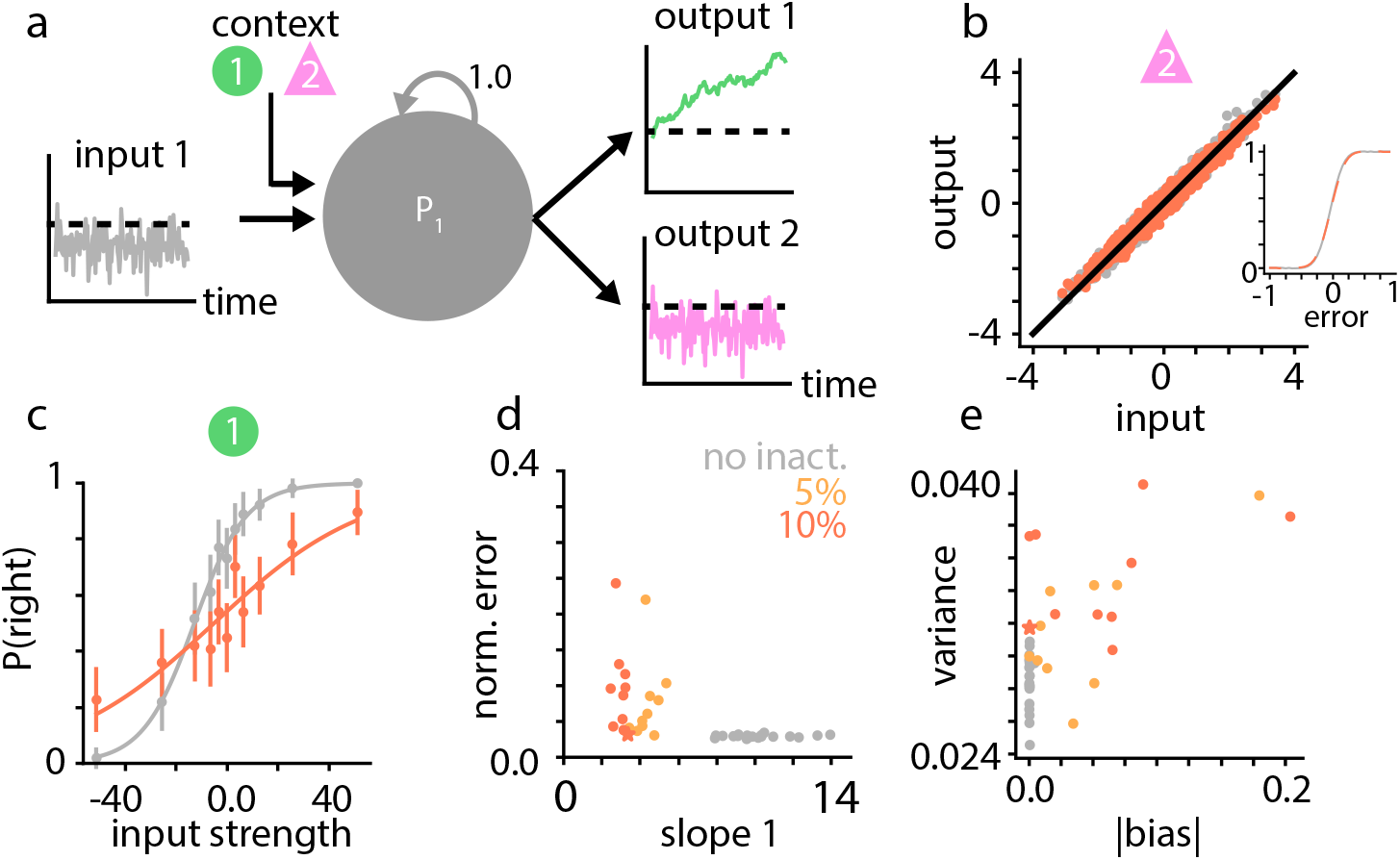
In a network trained to perform two different computations, inactivation can differentially affect performance. **a)** Schematic for a network integrating its input on 50% of trials (green), and reproducing the input on the remaining 50% of trials (pink). **b)** Plot showing the network input and output in context 2 against the identity (black dashed line). Grey indicates the pre-inactivation network, orange indicates the network after a 10% inactivation. Inset: cumulative density of errors across time and trials before and after inactivation. **c)** Psychometric function for responses in context 1 for input 1 before (grey) and after (orange) inactivation. **d)** Plot of the psychometric sensitivity in context 1 against the network output error (normalized by the variance of the sensory inputs) in context two for the pre-inactivation networks (grey), networks after a 5% inactivation (yellow), and after a 10% inactivation (orange). **e)** Plot of the bias and variance of the network output in context 2. Bars indicate ± 1 s.e.m. Starred points indicate the example network shown in panels b-c.

### 3.6 Short periods of relearning can compensate for inactivation

A key hidden assumption in our simulated experiments in the previous sections is that no additional task-specific learning can happen prior to testing the effects of the manipulation on behavior. This assumption is unlikely to be completely true, as plasticity and reinforcement mechanisms may continue to operate. In fact, cortical inactivation studies show many behaviors are only temporarily affected following the inactivation (Newsome & Pare 1988, Murray & Baxter 2006, Schiller et al. 1979, Rudolph & Pasternak 1999), a clear illustration of the brain’s remarkable capacity for learning through re-organization of its circuits. Similar recoveries may be expected in less severe experimental manipulations in which neurons are transiently inactivated, but the extent of additional learning required for adaptation to occur is less clear. To investigate the capacity of networks to adapt to inactivation and regain their performance through further task specific learning, we modified our model to allow network connections to continue to change at test time and investigated two biologically relevant variants of inactivation: long lasting and intermittent.

The first type of inactivation is implemented as a long-lasting disabling of the involved neurons, as used for all the previous sections. It is intended as an analogue of experimental manipulations using muscimol or other pharmacological agents, or designer receptors exclusively activated by designer drugs (DREADDs) (Wiegert et al. 2017), which affect target circuits for many minutes to days. In these cases, since the inactivation lasts for the majority of an experimental session (or multiple experimental sessions), circuits could eventually learn to compensate for the perturbation with sufficient additional task experience. What is remarkable in the context of the model is how little additional training is required.

Depending on the extent of manipulation, a few hundreds of trials were sufficient to compensate for the inactivation, much fewer than the number of trials required for the initial training of the network. Fig. 7a shows an example run where inactivation of 30% of the integrating population transiently caused the network to initially perform as poorly as it did before learning. To describe the trajectory of re-learning across networks, we measured the percentage of correct responses as a function of the number of retraining trials, for the same size of inactivation (30%). We found that the circuit robustly reached pre-inactivation performance with fewer than 500 retraining trials (Fig. 7b). This return to pre-inactivation performance was also mimicked in the underlying bi-stable attractor, with the bifurcation criterion *α* returning to positive values on the same time scale (Fig. 7c), indicating that the network has reconstructed its shallow bi-stable attractor.

**Figure 7:**
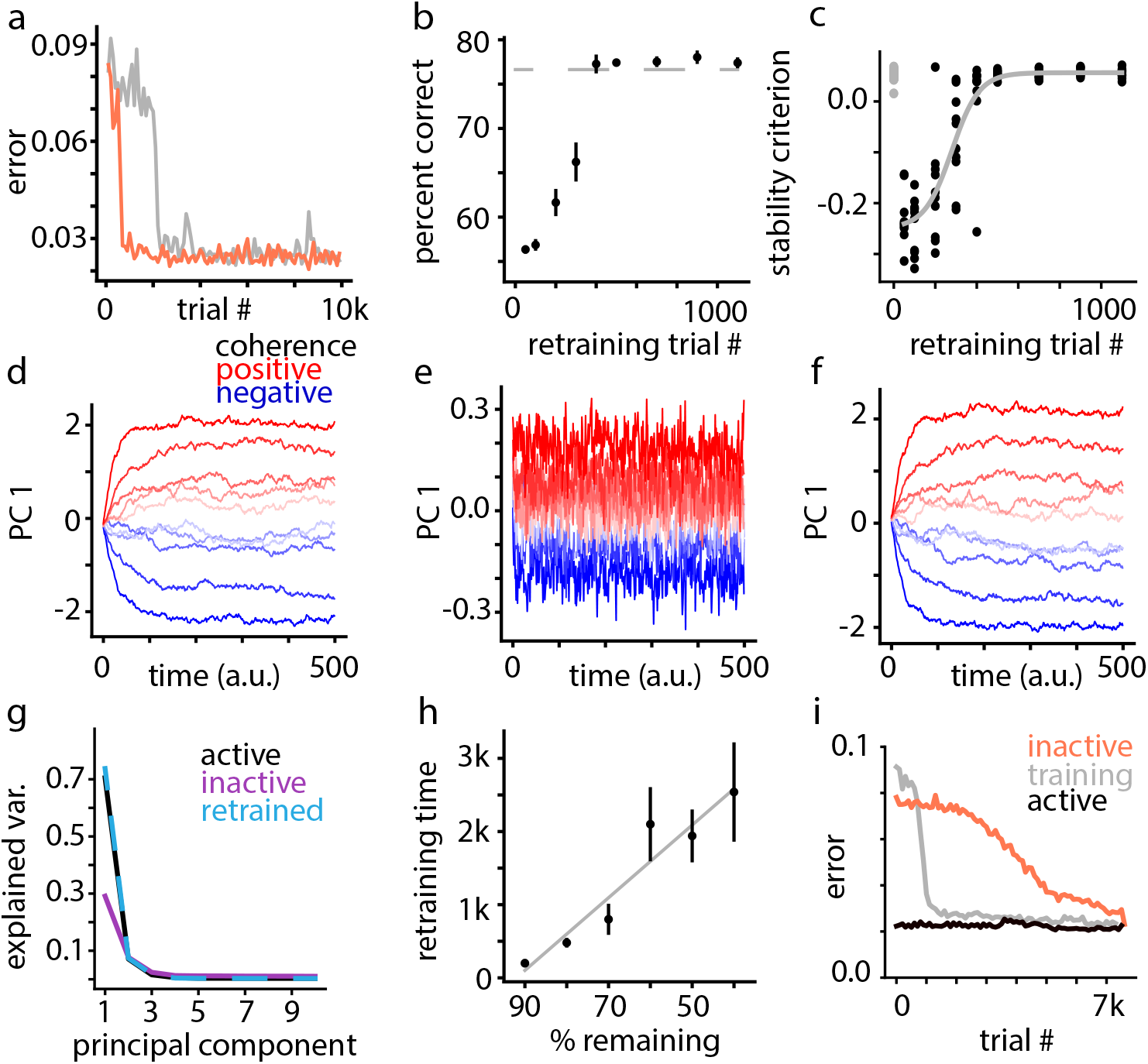
Perturbed networks can learn to compensate for inactivation but the speed of recovery depends on the timescale of inactivation. **a)** Post-inactivation training can be much faster than the initial training. The lines show changes in the MSE error of the hierarchical network of Fig. 1a during the initial training (gray) and during training after inactivation of 30% of P2 neurons. **b)** Percent correct as a function of retraining trials. The gray dashed line indicates the average network performance after initial training. **c)** The stability criterion as a function of the number of retraining trials. Data points show 10 different networks before inactivation (gray) and after inactivation of 30% of P2 neurons (black). **d)** Activity projected onto the first principal component for the network prior to inactivation. **e)** Activity projected onto the first principal component for the network after inactivation of 30% of P2 neurons. **f)** Activity projected on the first principal component for the network after post-inactivation retraining. **g)** Fraction explained variance as a function of the number of latent dimensions of the network responses after learning (black), after inactivation (purple), and after retraining (blue dashes). **h)** Mean retraining time as a function of the percentage of unperturbed neurons in the network. **i)** Retraining slows down considerably if inactivation of neurons occurs on a fast timescale that allows mixing perturbed and unperturbed trials during retraining. In this simulation, inactivation was limited to a random half of training trials. Integration error goes down slowly for perturbed trials (orange); here, trial # indicates only perturbed trials, not interleaved unperturbed trials. For the unperturbed trials (black), integration error remains close to the levels following the initial training (gray).

To directly visualize the impact of inactivation and relearning along the first axis of network variance, we compared the projection of the network activity onto its first principal component at the end of training (Fig. 7d), just after inactivation (Fig. 7e), and after relearning (Fig 7f; PCA performed separately on the neural activity at different time points). These show that the integration properties of the network largely collapse (although not completely) immediately after inactivation, and are fully and quickly restored by relearning. The relearning speed — time to reach virtually the same accuracy as the pre-inactivation network (within 0.5%) — is strongly correlated with the extent of inactivation: the larger the inactivated population, the longer it takes for function to be recovered by retraining (Fig. 7h). In our simulated network, inactivations as large as up to 30% of the neurons in the population still exhibited significantly faster relearning compared to the initial training time. This suggests there may be cases where compensation happens on the scale of one or a few sessions, similar to what an experimenter may use to assess the effects of the manipulation on behavior, potentially confounding these results.

The advent of optogenetics allows controlling the activity of neurons with millisecond resolution, leading to new experiments which interleave perturbed trials with unperturbed ones. These techniques are commonly considered the gold standard for causal manipulations as they offer millisecond temporal precision and enable targeting specific cell types. This improved precision and specificity is quite beneficial but does not remove the re-learning challenge mentioned above. The intact vs. the inactivated network can be thought of as two distinct dynamic states of the circuit. Repeated inactivation of largely the same group of neurons in a circuit, as in most optogenetic experiments, can provide opportunity for compensation even when inactivation is infrequent. Biological circuits could learn to use the silence of inactivated neurons as a contextual cue to switch behavior, or could redirect the computation in both states to the neurons that are not being directly manipulated.

To model an intermittent inactivation scenario similar to optogenetic manipulation experiments, we inactivated the network on a random subset (50%) of training trials, instead of tonically inactivating all neurons throughout retraining. In alignment to general intuitions that adaptation is less likely during transient inactivation, we found that it takes the network more inactivation trials to re-learn when inactivation is transient and infrequent (Fig. 7i). When neurons are only inactivated on 50% of retraining trials, it takes our network longer than its initial training time to compensate. This implies that transient inactivation techniques are likely more effective against inactivation-induced adaptation in biological networks, although compensation is still possible.

A possible criticism when interpreting these re-learning results is that the optimization procedure used for learning is not biologically realistic, and that the dynamics of re-learning might look very different when the network connections adapt via local synaptic plasticity rules. To assess the generality of our results, we trained the network using Random-Feedback Online Learning (RFLO), a biologically-plausible alternative to backpropagation through time (Murray 2019). We also replaced mean-squared error with a loss based on binary decision outcomes, as a more realistic feedback signal to the network (see Methods). In Suppl. Fig. S3b-g we repeat the training and inactivation experiments for different inactivation sizes in this new model. We find that the qualitative features of the network solution and the post-inactivation loss of function match those shown in Fig. 2. In particular, the network is learning bounded integration (Fig. S3a), with a moderate loss-of-function in the psychometric and chronometric functions for a 40% inactivation, and near-total loss-of-function for a 75% inactivation. As the network continues to learn after inactivation, it is able to restore its mean decision time and performance much more rapidly than the original training time for both the 40% (Fig. S3f,h) and 75% (Fig. S3g,i) inactivations, suggesting that local synaptic plasticity can also support fast recovery of function after partial circuit inactivation.

## 4 Discussion

A main quest of neuroscience is to identify the neural circuits and computations that underlie cognition and behavior. A common approach to achieve this goal is to first use correlational studies (e.g., electrophysiological recordings) to identify the circuits whose activity co-fluctuates with task variables (e.g., stimulus or choice), and then perturb those circuits one by one as subjects perform the task. Loss of task accuracy following lesions or transient inactivation of a circuit is commonly interpreted as evidence that the circuit is “necessary” for the underlying computations. The converse, however, need not be true. Of course, if the inactivated circuit is not involved, the behavior remains unaffected. But negative results can also arise because of other reasons, which challenge the impulse of embracing the null hypothesis.

The conclusion that negative results in perturbation experiments are not readily interpretable is not new. In fact, it is common knowledge in statistics that inability to reject a null hypothesis (e.g., circuit X is not involved in function Y) is not evidence that the null hypothesis is correct, especially if the true causes of the results remain unexplored and not included in hypothesis testing. In practice, however, there is a growing abundance of publications that interpret negative results of perturbation experiments as lack of involvement of a circuit in a mental function. In many cases, experimenters perceive their negative results as important because they seem to contradict established theories (e.g., role of posterior parietal cortex in perceptual decisions (Katz et al. 2016)). A popular approach is to make negative results palatable or interpretable through establishing a “dissociation” by showing that the same perturbation in a different task or a different circuit yields positive results. Aside from statistical and logical concerns about the validity of this remedy (Yoshihara & Yoshihara 2018, Dunn 2003), our results reveal key challenges often ignored in the interpretation of negative results: they can emerge from robustness to perturbation due to the architecture of the affected circuit or the bigger network that the circuit belongs too; circuits may continue to learn, at a much faster scale than we tend to expect, even when experimentalists don’t wish it); and circuits that perform multiple tasks could have developed solutions that are robust to perturbation effects in one task and sensitive in another task, even when the circuit is involved in both.

Our results emphasize the need for further exploration following negative results and point at exciting directions for followup experiments. There is already evidence that circuits adapt to both the transient inactivations (Jeurissen et al. 2021, Fetsch et al. 2018) and permanent lesions—for example the brain’s impressive robustness to extensive and gradual dopamine neuron loss in Parkinson’s disease (Zigmond et al. 1990). However, we do not know which brain circuits adapt to perturbations, or under what experimental conditions. Answering this key question would be aided if experiments document behavior from the very first administration of the perturbation protocols and devise methods to quantify different learning opportunities inside and outside of the experimental context. It is also essential to know about the larger network engaged in a task and the parallel pathways that could mediate behavior. Association cortex, for example, often includes recurrent and diverse connections, which give rise to many possible parallel pathways. Such complex networks often prevent straightforward conclusions from a simple experimental approach that perturbs a single region in the network. More elaborate experimental designs and carefully developed computational models aid overcoming this complexity. We recommend quantification of behavior not be limited to choice (a discrete measure) and include more sensitive, analog measures, such as reaction time (Results 3.4), which was affected in our distributed network inactivation simulations, even when accuracy was seemingly unaffected. We also see strong advantage in augmenting single region perturbations with simultaneous perturbation of a collection of network nodes, chosen based on network models. Another valuable approach is to simultaneously record unperturbed network nodes to quantify the effects of perturbation on brain-wide response dynamics and identify adaptive mechanisms that could rescue behavior (Li et al. 2016). From our perspective, negative behavioral results in a perturbation experiment are not the end point of the experiment. Rather, they are just a step toward a deeper understanding of the neural mechanisms that shape the behavior.

Our models in this paper focus on a well-studied perceptual decision-making task: direction discrimination with random dots. Understanding the dynamical mechanisms of computation in our circuits proved necessary for understanding their response to inactivation. Trained networks implement an approximation of the drift-diffusion model, and exhibit low-dimensional integration dynamics in response to sensory stimuli. We characterized the learned solutions by their phase portrait properties, and found that networks approximated sensory integration using a shallow bistable attractor (Wong & Wang 2006, Strogatz 2018), whose disruption was closely correlated with loss-of-function in inactivated networks; further, relearning reconstructed a bistable attractor within the network. Our analysis shows that what matters in a circuit, irrespective of the status of individual neurons, is the integrity of its computational structure. This makes statistical approaches that aim to extract this structure directly from measured population responses particularly valuable (Zhao & Park 2016, Nassar et al. 2018, Duncker et al. 2019).

Though different tasks and network structures will likely lead to variations in the nature of learned solutions and responses to inactivation, in many respects the random dots motion discrimination task and our neural network architectures serve as a microcosm of a more general phenomenon, which encompasses both artificial and biological systems. Results in the recurrent neural network literature have shown that significant variations in the response properties of individual network units (vanilla RNNs, GRUs, or LSTMs) tend to produce similar canonical solutions to simple decision-making tasks, embedded in a low-dimensional subspace of the network’s dynamics (Maheswaranathan et al. 2019). The exact mechanism of loss-of-function in response to inactivation may differ, but our expectation is that independent of architecture, any trained system–including those used by biological systems–will learn to approximate the optimal canonical computations (here, integration) required for the task. We examined only the simplest possible implementation of parallelism in our network architecture, and showed that even this was sufficient to greatly increase the system’s robustness to neural inactivation. Real neural circuits likely show this phenomenon on a much larger scale by virtue of involving far more neurons, with more natural redundancy.

The particular learning algorithm that drives the organization of the circuit is not important for the observed effects: a biologically-plausible learning algorithm (Murray 2019), with more realistic feedback, learned a similar computational structure, and showed similar inactivation and relearning effects to brute-force optimization via backpropagation through time. It is likely that *any* algorithm that closely aligns with gradient descent on a task-specific loss will produce similar effects.

Though we found that the simplest network architectures were in general not robust to inactivations, many basic architectural modifications could produce resistance to perturbations. In particular, redundancy of function (two parallel, independent attractors) embedded within the network produced inactivation resistance; this suggests that multi-area recordings are important for assessing whether the effects of inactivation have been compensated for by another neural population. Further, in our distributed circuit, inactivation still affected reaction time, suggesting that reaction time or other analog aspects of behavior may be more sensitive than choice to inactivation effects.

As our results show, when multiple tasks are used to assess the effects of inactivation, interpreting situations that do not affect one task but do affect another task can be difficult, especially if the two tasks involve different computations. We have shown such a situation, where a modeled neural circuit is known to contribute to both tasks, but perturbations have a large effect on one task, and a small effect on the other. Recapitulating previous empirical results (Mante et al. 2013), we found that a single recurrent network could form multiple functional units for solving separate tasks, depending on a contextual signal. Whether lesions to a random subset of neurons affected both tasks depended strongly on which tasks were being performed. In particular, we found that a network performing integration on two separate sensory streams suffered similar performance impairment across both tasks, but that a network performing both an integration task and a replication task only lost function on the integration task. This implies that more generally, if the different computations implemented by a circuit have different forms (e.g. one task requires a bistable attractor and one does not), inactivations can differentially impact function across the two tasks, because one task is either more complex, or less robust to inactivation.

Furthermore, we found that even if neural circuits play a direct causal role in a computation, loss-of-function in response to inactivation of a subset of neurons can be transient. Longer-term inactivation of neurons in our circuit, in the presence of active learning, allowed networks to rapidly compensate. This compensation occurred on a time scale much faster than the original training time, likely because inactivation does not completely destroy the network’s previously learned computations. Recovery of behavior has been observed before in experiments (Murray & Baxter 2006, Newsome & Pare 1988, Fetsch et al. 2018, Rudolph & Pasternak 1999, Jeurissen et al. 2021). Overall, drawing conclusions about the causal role a circuit plays in a given computation can be difficult without first analyzing the transient responses of animals immediately after inactivation — a commonly omitted or poorly documented aspect in many studies.

Fast time-scale inactivation techniques (e.g., optogenetics) (De et al. 2020, Luo et al. 2018, Wiegert et al. 2017, Afraz et al. 2015) have greatly increased in popularity as they allow precise control of the affected neurons with sub-second resolution. As we show here, brief periods of inactivity interspersed with normal activity also make it harder for a learning system to identify and adapt to the perturbation. However, compensation can occur even for fast optogenetic perturbations (Fig. 7), as has been observed experimentally (Fetsch et al. 2018). But such compensations tend to take longer compared to techniques in which the inactivation is more sustained (Fig. 7). This longer adaptation may be a result of destruc-tive interference during re-learning, where the synaptic changes needed to improve performance during perturbation are misaligned and cancel those in the absence of perturbation, thus slowing down learning overall. For our example direction-discrimination task, once compensation does occur, it could take the form of two separate attractors, one dedicated to performing during the perturbed condition, and the other for the unperturbed condition. Alternatively, the network may converge to a single attractor, modified from its original solution such that does not include the inactivated subset of neurons.

The phenomena observed in this paper that can produce negative results in an inactivation study— redundancy, rapid relearning, differential sensitivity to inactivation across tasks—are all problems more general than just evidence integration (Vaidya et al. 2019, Wolff & Ölveczky 2018). Here we have provided several suggestions for identifying the effects of causal manipulations in neural circuits, and have provided several cautionary tales based on the choice of a particular architecture and task. Circuits before and after manipulation are only tenuously related, and drawing conclusions about the function of natural circuits from the effects of inactivation can be quite difficult. Implementing proper controls for these effects and applying careful interpretations of observed experimental results in terms of the system’s computational structure will benefit inactivation studies across a breadth of subfields.

## 5 Methods

Cognitive functions depend on interactions of neurons in large, recurrent networks. To explore the utility and limitations of inactivation and lesion studies for discovering the flow of information and causal interactions in these networks, we simulated recurrent neural network (RNN) models with different degrees of complexity and selectively inactivated sub-populations of neurons within the simulated networks. The models were trained to perform simple perceptual decisions, commonly used for investigating cortical and subcortical neural responses and their causal contributions to behavior (Katz et al. 2016, Fetsch et al. 2018, Zhou & Freedman 2019b, Hanks et al. 2015). Our simulations and theoretical exploration focus on the direction discrimination task with random dots (Newsome & Pare 1988, Roitman & Shadlen 2002) as a canonical example of perceptual decision-making tasks.

### 5.1 Implementation of RNNs

We simulated an RNN performing a random dots task. To ensure convergence to an optimal set of weight parameters, we trained the RNNs in PyTorch (Paszke et al. 2019) using backpropagation through time (BPTT) and Adam (Kingma & Ba 2014) with a learning rate of 2 × 10^*−*6^. Each network was trained over 25,000 trials and tested on a separate group of 1500 trials for investigating network computations, task performance, and susceptibility to various activity perturbations. The time steps can be mapped to physical units of time (e.g., 10 milliseconds), but we avoid doing so as our conclusions are invariant to the exact definition of time steps. A univariate input sampled from a Gaussian distribution, *s* ∼ N(*kC*, 1), was applied at each time step. *C* is the motion strength (coherence), and *k* is a sensitivity parameter that translates motion strength to sensory evidence. The variance of the input evidence to the network was set to 1. In our simulations, the sensitivity was *k* = 0.4, and *C* was randomly drawn on each trial from a discrete set: [-0.512, -0.256, -0.128, -0.064, -0.032, 0, 0.032, 0.064, 0.128, 0.256, 0.512]. Positive and negative motion strengths indicate rightward and leftward directions, respectively. The network was trained to discriminate the two motion directions based on input evidence, as explained below.

Independent normal noise (*η*) was injected into each neuron at each time step with variance 0.01. These variables combined give the following update equation for the RNN:

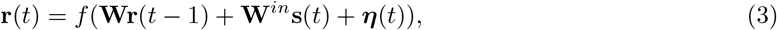

where **W** is the recurrent weight matrix, **W**^*in*^ is the input weight matrix, and *f* (·) is the tanh nonlinearity. The network output is given by:

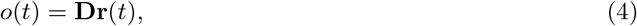

where **D** is a 1 × *N* linear decoder, and *N* is the number of neurons in the network.

We trained the network to integrate these inputs through time, setting our loss to the mean-squared error (MSE) between the network output and an integrated decision variable (*DV*) given by:

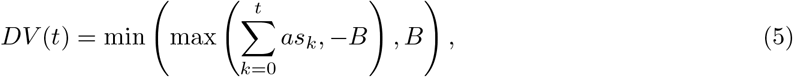

with proportionality constant *a* = 0.025 to keep the integrated variable within the dynamic range of the RNN, and *B* = 0.5 giving the bounds on integration. This results in the loss ℒ is given by:

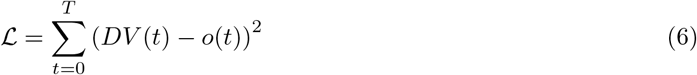

The nonlinearity in Eq. 5 limits the dynamic range of the DV and ensures accurate representation of low-magnitude DVs in the small pool of neurons in the RNN model. Since the majority of motion strengths in our simulations are weak, accurate representation of low-magnitude DVs are crucial for task performance.

During training, the length of each trial was selected randomly from an exponential distribution, *T*∼ 100 + exprand(200), with a maximum duration of 500 time steps. During testing, we used the maximum stimulus duration to ensure all trials terminated by reaching the decision bounds, enabling us to determine both choice and decision time on each trial.

### 5.2 Simple circuits

To begin, we trained a two-population network, where the first population (*P*_1_) receives the evidence input, *s*(*t*), and a linear decoder (Eq. 4) reads out the integrated input from the second population (*P*2). Connections between the two populations are feedforward, with each feedforward connection having a probability of 0.3 of being nonzero, but connections within each population are all-to-all, as shown in Fig. 1a For the sake of simplicity, there were no feedback connections from *P*_2_ to *P*_1_ (connection probabilities are shown in Fig. 1a). The first population had 30 neurons, and the second had 60. These two populations fulfill the roles of a low-level sensory population that relays input information, and a higher-order population that integrates the information for a decision. This network is much simpler than the circuit that underlies sensory decisions in a mammalian brain, where motion selective sensory neurons (e.g., MT neurons in the primate brain) pass information about the sensory stimulus (Newsome & Pare 1988, Salzman et al. 1990, Britten et al. 1992) to a large network of association cortex, motor cortex, and subcortical areas that form the decision (Horwitz & Newsome 1999, Ding & Gold 2013, Roitman & Shadlen 2002, Mante et al. 2013, Kiani, Corthell & Shadlen 2014, Kim & Shadlen 1999). However, our simple circuit lends itself to mathematical analysis, can be trained without adding structural complications, and can be used for systematic exploration of inactivation effects.

To emulate lesion or inactivation experiments, we selectively inactivated a fixed group of neurons in the network. In lesion or long-term inactivation experiments (e.g., muscimol injections or DREADS) the connections remained affected throughout all trials in a testing block. In inactivation experiments with fast timescales (e.g., optogenetic perturbations), the connections were affected for a random subset of trials intermixed with other trials in which all connections and neurons functioned normally.

We systematically varied the proportion of affected neurons in the population in distinct simulations. A weak perturbation affected 5-10% of neurons, a medium-strength perturbation 20% of neurons, and a strong perturbation 30% of neurons (Fig.1a).

### 5.3 Complex circuits

In the distributed network that subserves perceptual decision-making, multiple circuits could operate in parallel, performing similar operations. To investigate the impact of this possibility on our inactivation results, we organized the network into two unconnected populations, each with 30 neurons. The connection probabilities are given in Fig. 4a. We totally inactivated one of the two sub-populations and analyzed the effect on network responses. Note that our simulation is not meant to capture the full complexity of the equivalent brain circuits but rather to offer a minimalist design that captures a key complexity commonly observed in brain networks: parallel processing of sensory information in a variety of frontal and parietal cortical regions (e.g., lateral intraparietal, frontal eye fields, and lateral and medial prefrontal areas of the monkey brain).

### 5.4 Analysis of neural responses

After training, we applied several analyses to characterize the nature of the network computations and effects of perturbations. Because there is not an explicit reaction time in our training framework, we set symmetric decision boundaries on the network output *o*(*t*) as a proxy for reaction time. We quantified the time until *o*(*t*) reached one of the boundaries on each trial. The crossed boundary dictated the choice on the trial and the time to bound determined the decision time. Formally, the reaction time is given by:

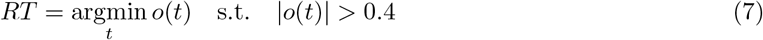

and the choice is given by:

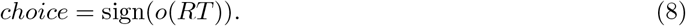

For each trained RNN, we constructed a psychometric function by measuring the proportion of ‘left’ and ‘right’ motion choices, and we fit the psychometric function using the following logistic regression:

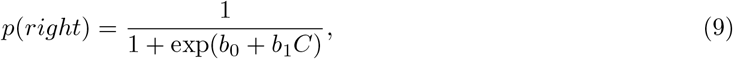

where *p*(*right*) is the proportion of ‘right’ choices, and *b*_*i*_ are regression coefficients. *b*_0_ reflects the choice bias, and *b*_1_ the sensitivity of choices to changes in motion strength.

We constructed chronometric functions (Fig. 2c) by stratifying the mean decision times as a function of motion strength. For simplicity, we fit the chronometric functions with nonlinear regression using the following bell-shaped function:

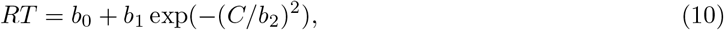

where *RT* is the network’s decision time.

To explore the dynamics of neural responses, we performed PCA on the network activity over time and trials. We found that the majority of population response fluctuations lie within a single dimension (See Fig. 1h). We analyzed neural trajectories associated with each choice by averaging neural firing rates within the output population for choices to the left, for each motion strength. A perfect integrator would have a mean response linearly increasing through time, and the slope of that linear increase would vary linearly with changes in motion strength (Shadlen et al. 2006). To verify this, we fit the mean output (*o*_*t*_) through time by linear regression (Fig. 1c), and plotted the slope of this fit as a function of coherence (Fig. 1d). Further, a perfect integrator would show a linear increase of variance over time. We measured the variability of the network responses at each time point and for each motion strength by quantifying the empirical variance of the network output across test trials with the same motion strength.

### 5.5 One-dimensional approximate dynamics and pitchfork bifurcation

Given the low dimensional structure of the trained RNN dynamics, we can provide a one-dimensional approximation of our RNN by projecting the network dynamics along its first principal component. Let **V** be eigenvectors corresponding to distinct eigenvalues of the covariance matrix of the network activity, obtained by combining trials across time and stimuli. These normalized vectors define an orthonormal basis, with the first and *n*-th axis corresponding to the direction of maximum and minimum variance, respectively. The activity of the network in this rotated coordinate system becomes **r**_*rot*_(*t*) = **V**^⊤^**r**(**t**). Using Eq. 3, this leads to dynamics:

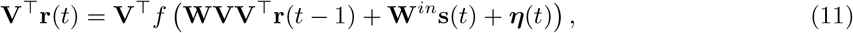

where we have used the fact that the basis is orthonormal, i.e. **VV**^⊤^ = **I**. Substituting our definition for **r**_*rot*_, we have:

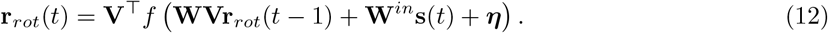

Focusing on the first dimension, along the axis of maximum variance, yields:

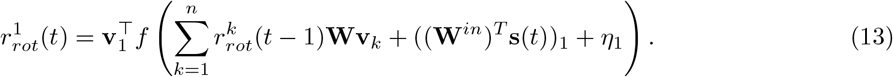

where **v**_*k*_ denotes the *k*-th eigenvector, and 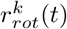 is the *k*-th entry of **r**_*rot*_(*t*). Assuming that the system is largely one-dimensional, the expression for the dynamics can be further simplified as:

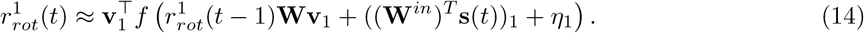

This approximation effectively discards the contribution of the remaining dimensions, under the assumption that their effect on the network dynamics is minimal, i.e. 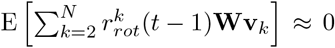, which holds empirically for our trained networks. In Fig. S2, we demonstrate for a trained network that the approximate dynamics closely match the true neural dynamics along its highest variance dimension 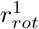.

Having derived a one-dimensional dynamical system approximation to the RNN activity, we can use phase-plane methods to determine the nature of the learned dynamics. We are interested in the geometry of the solution our network finds, which leads us to assess its fixed point dynamics in the absence of input and noise (**s**(*t*) = 0, *η*_1_(*t*) = 0). Finding these fixed points involves finding the solutions of equation:

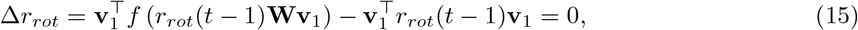

where Δ*r*_*rot*_ = *r*_*rot*_(*t*) − *r*_*rot*_(*t* − 1), and we have used the fact that the eigenvectors are normalized, i.e.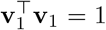. Further, using a Taylor approximation of tanh about 0, *f* (*x*) ≈ *x*−*x*^3^, and rearranging the terms simplifies the equation to:

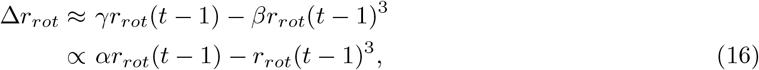

where 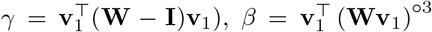 is empirically positive, *α* = *γ/β*, and (·)^°3^ denotes an element-wise cube. The resulting equation is cubic, meaning its fixed point equation (Δ*r*(*t*) = 0) has up to 3 solutions. This generally results in a topology with two stable fixed points separated by one unstable fixed point. These points coalesce into a single stable fixed point, *r*_*rot*_(*t*) = 0, when the coefficient of *r*_*rot*_(*t* −1) changes from positive to negative, with the system undergoing a *supercritical pitchfork bifurcation* (Strogatz 2018).

For the network to work properly, it needs to be in the regime with two stable attractors, with an abrupt degradation once reaching the critical point for the phase transition. For our approximate dynamics, this transition occurs once:

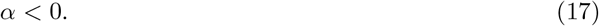

For this reason, we refer to the parameter *α* as the bifurcation criterion.

### 5.6 Re-learning with feedback after perturbation

To investigate whether perturbations have a lasting impact on the performance of a network with plastic neurons, we permitted the simplified hierarchical network in Fig. 1a to be trained following inactivation. Two training regimes were used to emulate different experimental techniques with slow and fast timescales for inactivation. In both regimes, we silenced a fraction of neurons in the P2 population and allowed the connection weights of the remaining neurons to change through relearning. The first retraining regime was designed to emulate lesion and pharmacological inactivation studies, which affect the circuit for an extended time, ranging from a whole experimental session to permanent. In this regime, the affected neurons remained inactive throughout the retraining period. The second regime was designed to emulate optogenetic or other techniques with faster timescales, which allow interleaving perturbations with unperturbed trials. In this regime, we silenced the affected neurons in a random half of retraining trials and allowed them to function in the other half; synapses were modified in all trials.

To assess the efficacy of retraining in restoring the network performance, we used the state of synapses at various times during retraining to simulate 1500 test trials and calculate the percentage of correct responses. Connection weights were kept constant in the test trials. Additionally, we calculated the projection of the network activation onto its first principal component following the initial training, after the inactivation and prior to retraining, and at various times during retraining. Finally, we calculated the stability criterion (Eq. 17).

### 5.7 Task-dependence of inactivation effects

To examine the effects of inactivation on networks performing multiple tasks, we trained a 1-population model (*N* = 100) (Fig. 1a) to perform contextual integration (Fig. 5a), where the network was required to integrate one of its two inputs depending on the context signal it received. We randomly selected a task as the main readout of the network to perform on each trial with 50% probability. Thus, the objective function for training was:

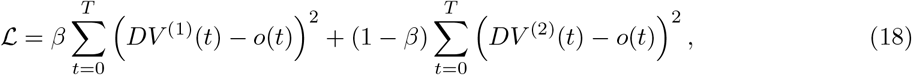

where *β* ∼ *Bernoulli*(0.5), and *DV* ^(1)^ and *DV* ^(2)^ are given by Eq. 5 for stimuli *s*^(1)^ and *s*^(2)^, respectively. To allow the network to properly infer context, we fed the binary cue *β* as a 1-hot feedforward input to the population ([*β* (1 − *β*)]). We trained the network on 300,000 trials, so that it reached good performance on both tasks. We then applied weak (%5 of neurons) and medium-strength (%10 of neurons) inactivation to the network. Here we redefined the ‘weak’ and ‘medium’ inactivation magnitudes because we applied the inactivation to an entire 100-neuron network, rather than selectively to one population.

We calculated the psychometric and chronometric functions, as well as the sensitivity and bias, as in Section 5.4, but we also calculated the psychometric function where stimuli were given by the off-context input instead of the on-context input. This allowed us to see if the off-context inputs affected network behavior in any way.

#### 5.7.1 Integration and replication

Alternatively, rather than integrate one of two inputs, we took the same network and fed in a single stimulus. Depending on the context, the network was either required to integrate the stimulus (*DV* ^(1)^ given by Eq. 5), or replicate the stimulus, with *DV* ^(2)^(*t*) = *s*(*t*).

### 5.8 Biologically plausible learning

It is highly unlikely that a neural system could receive detailed feedback about the difference between a decision variable and an integrated target trajectory. There is no supervised signal for this target trajectory, and if a neural system was able to construct the target, why not use it to solve the task instead? Instead, an animal is much more likely to use reward feedback that it receives about its classification. To verify that our results hold in this situation, we adapted our training to use a cross-entropy loss function:

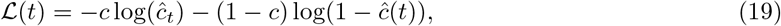

where *c* = sign(*C*), and *ĉ* = *σ*(*o*(*t*)), where *σ*(·) is a sigmoid nonlinearity.

For the simulations here, we evaluated the loss at every time step, though we achieved qualitatively similar results with only end-of-trial evaluation.

BPTT is well-established as a biologically implausible learning algorithm (Werbos 1990). Many studies have constructed approximations or alternative formulations of BPTT that are biologically plausible, but these algorithms often have different stability properties or biases (Marschall et al. 2020), or are not guaranteed convergence to the same solution (or convergence at all). To verify that our results still hold for a biologically plausible learning algorithm, we selected RFLO (Murray 2019), where recurrent weight updates are given by:

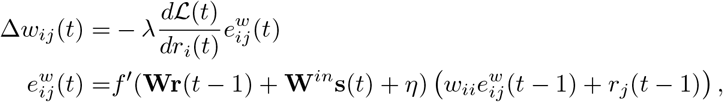

where *λ* = 0.001 is the learning rate, and where the second equation is an ‘eligibility trace’, which is updated continuously at each synapse, and requires only information available at the pre- and postsynapse.

This has the form of a three-factor plasticity rule (Frémaux & Gerstner 2016), where a reward signal 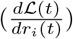 is fed back and combined with pre-synaptic and post-synaptic Hebbian coactivation to produce the weight update. In our case, we allowed the feedback weights to be given by direct differentiation of the objective function, but for added biological realism, these weights could be learned (Akrout et al. 2019) or random (Murray 2019, Lillicrap et al. 2016) and still achieve good performance.

The updates for the input weights are analogous:

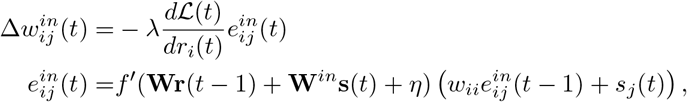

and the updates for the decoder are simply given by:

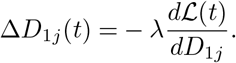

For the sake of computational efficiency, for these simulations we decreased the number of time steps to 30 steps per trial with a fixed duration (10,000 trials), and increased the signal-to-noise ratio of individual stimuli by taking *s* ∼ 𝒩 (*kC*, 0.1) for *k* = 0.4. Further, we gave our network only one population of neurons (*N* = 60) with all-to-all connectivity. We also trained our networks with a larger amount of intrinsic noise (*σ* = 0.6) to verify that our results hold for noisier neurons.

Because these simulations had modified parameters and a different objective function, we had to reset our decision threshold to achieve qualitatively similar psychometric functions. We set the threshold for decisions in these simulations to 1: we arrived at this value by requiring near-perfect choice accuracy for strong coherence stimuli, and a chronometric function whose mean response times peaks at 0 coherence. These features clearly need not be achievable for *any* coherence if the task has not been well-learned, but we found in practice that a threshold value of 1 gives psychometric and chronometric functions similar to experimental data.

Because our networks had a different number of neurons and a different objective function, we also recalibrated the magnitudes of our inactivations. Our ‘weak’ inactivation in these simulations targeted 40% of neurons, and our ‘strong’ inactivation targeted 75% of neurons. The method of performing our inactivation was identical to the previous section.

## Acknowledgements

We thank Michael Shadlen, Jean-Paul Noel, Saleh Esteki, Gouki Okazawa, Michael Waskom, John Sakon, Danique Jeurissen and S. Shushruth for inspiring discussions and feedback on earlier versions of the manuscript. We thank Owen Marschall for sharing code to implement the RFLO algorithm. This work was supported by the Simons Collaboration on the Global Brain (542997), National Institute of Mental Health (R01 MH109180), and the Alfred P. Sloan foundation. Additionally, RK was supported by a Pew Scholarship in the Biomedical Sciences, and a McKnight Scholar award. CS was supported by National Institute of Mental Health (1R01MH125571-01), the National Science Foundation (NSF Award No.1922658) and a Google faculty research award.

---

**Figure S1:**
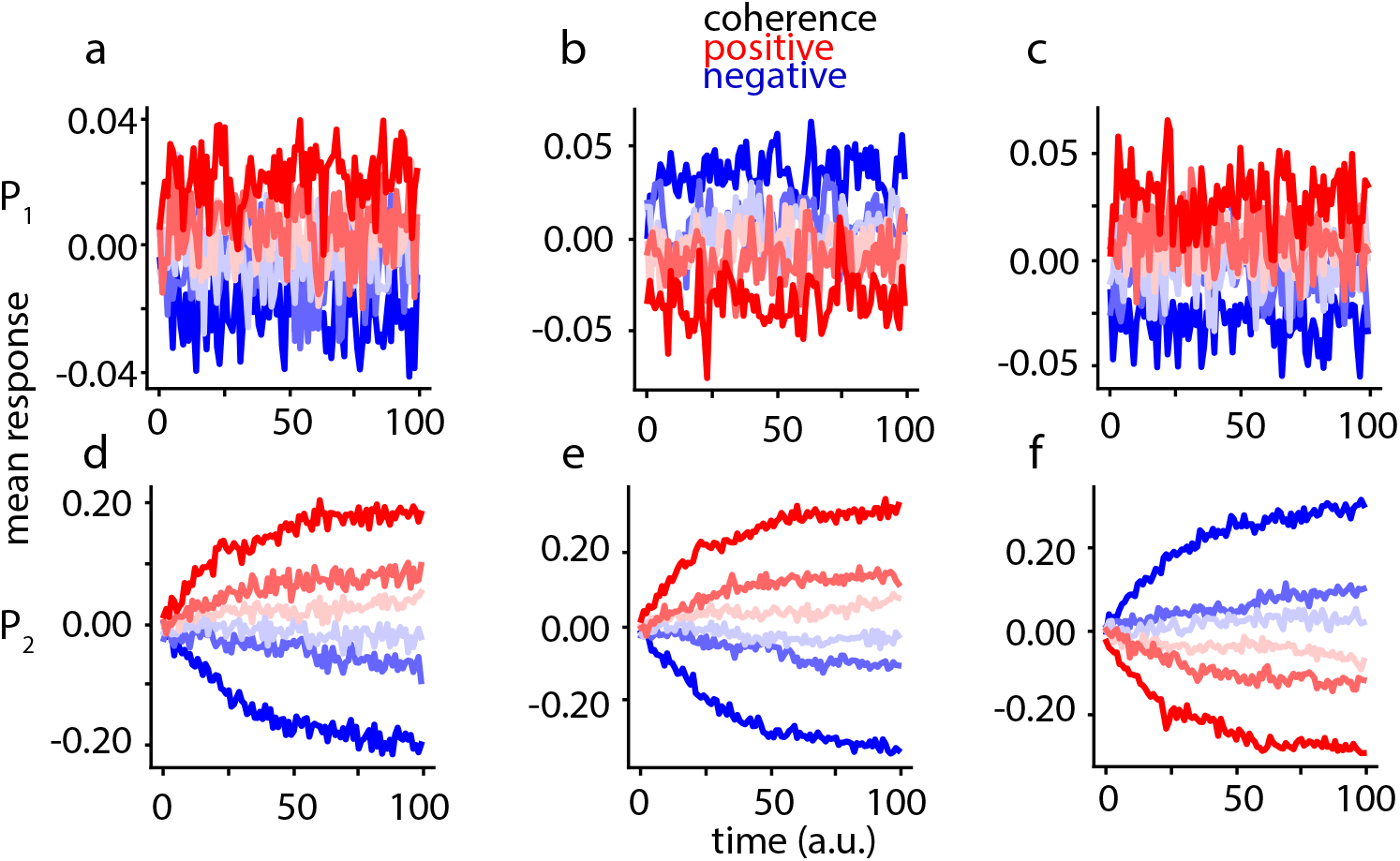
Example neuron response profiles from the simple integration network. **a-c)** Example stimulus-conditioned mean neuron responses from P1. **d-f)** Example stimulus-conditioned mean neuron responses from P2. Colors indicate the coherence strength and sign (blue indicates leftward motion, red indicates rightward motion).

**Figure S2:**
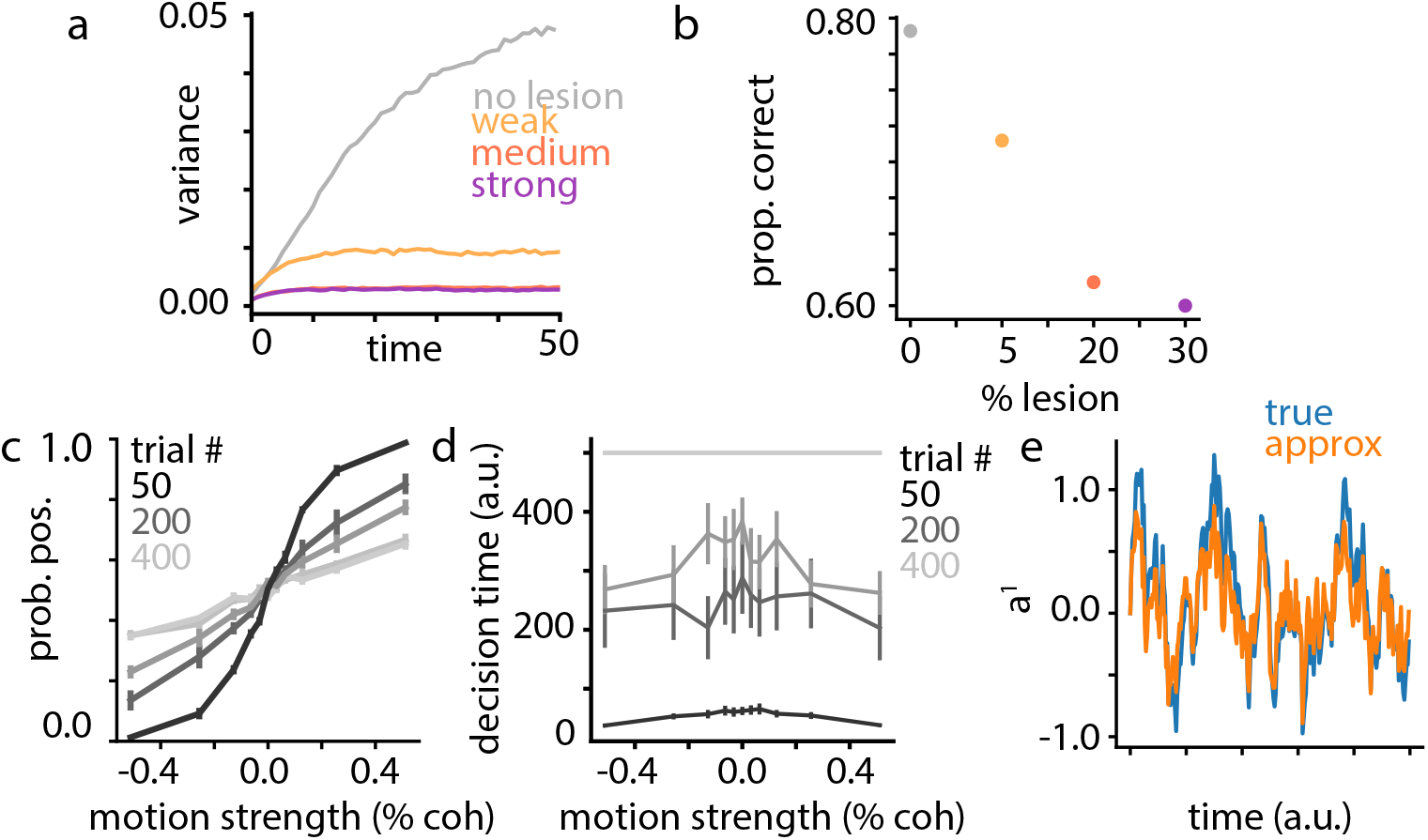
Additional analysis on the effects of inactivation and relearning. **a)** Variance of the network output through time as a function of the perturbation magnitude **b)** Proportion of incorrect choices as a function of the number of inactivated neurons in the output population. **c)** Psychometric function as a function of the number of retraining trials given to the network for a strong inactivation **d)** Same as (c), but for the chronometric function. Bars indicate ± 1 s.e.m. across 10 simulated networks **e)** Difference between the true (Eq. 14) and approximate (Eq. 16) dynamics for a trained network over 1 trial projected onto the 1st principal component.

**Figure S3:**
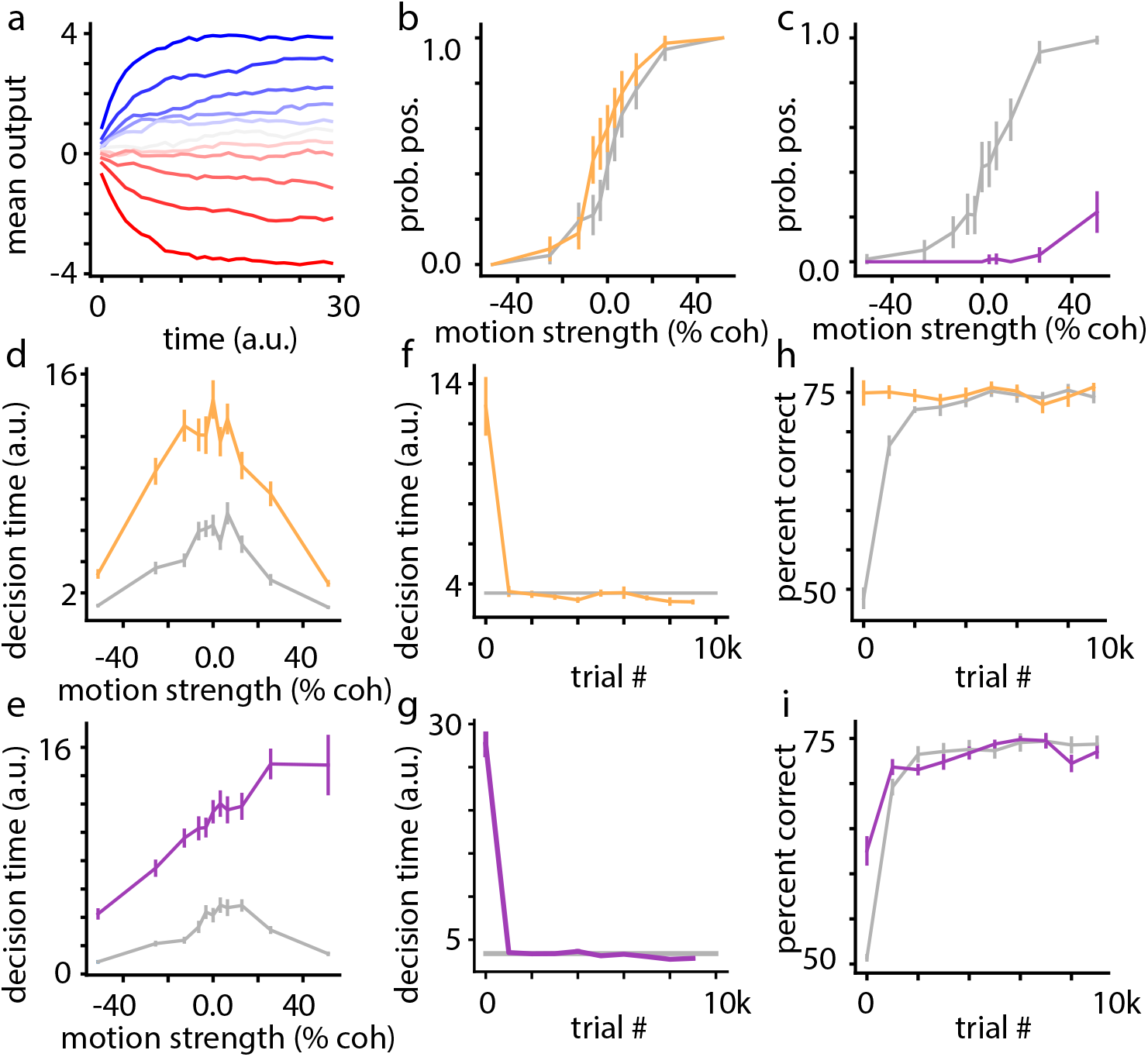
Inactivation and relearning analysis for a network trained with biologically-plausible learning. **a)** Mean output throughout time, stratified by coherence. **b)** Psychometric function after training (grey), and after a 40% inactivation (yellow). **c)** Same as (b), but for a 75% inactivation (purple). **d)** Chronometric function after training (grey) and after a 40% inactivation (yellow). **e)** Same as (d), but for a 75% inactivation (purple). **f)** Mean decision time for a 40% inactivation throughout retraining (yellow), compared to the asymptotic mean decision time prior to inactivation (grey). **g)** Same as (f), but for a 75% inactivation (purple). **h)** Percent correct for a 40% inactivation throughout retraining (yellow), compared to the percent correct through the original training. **i)** Same as (h), but for a 75% inactivation (purple). Bars indicate ±1 s.e.m. across 1000 test trials for (b), (c), (d), and (e), and ±1 s.e.m. across 10 simulated networks for (f), (g), (h), and (i).

